# pathwayPCA: an R package for integrative pathway analysis with modern PCA methodology and gene selection

**DOI:** 10.1101/615435

**Authors:** Gabriel J. Odom, Yuguang Ban, Lizhong Liu, Xiaodian Sun, Alexander R. Pico, Bing Zhang, Lily Wang, Xi Chen

**Affiliations:** Division of Biostatistics, Department of Public Health Sciences, University of Miami, Miller School of Medicine, Miami, FL 33136, USA; Sylvester Comprehensive Cancer Center, University of Miami, Miller School of Medicine, Miami, FL 33136, USA; Dr. John T. Macdonald Foundation Department of Human Genetics, University of Miami, Miller School of Medicine, Miami, FL 33136, USA; Institute for Data Science and Biotechnology, Gladstone Institute, San Francisco, CA 94131, USA; Department of Molecular and Human Genetics, Baylor College of Medicine, Houston TX 77030, USA

**Author notes:** To whom correspondence should be addressed. Xi Chen; Lily Wang.

## Abstract

With the advance in high-throughput technology for molecular assays, multi-omics datasets have become increasingly available. However, most currently available pathway analysis software provide little or no functionalities for analyzing multiple types of -omics data simultaneously. In addition, most tools do not provide sample-specific estimates of pathway activities, which are important for precision medicine. To address these challenges, we present `pathwayPCA`, a unique R package for integrative pathway analysis that utilizes modern statistical methodology including supervised PCA and adaptive elastic-net PCA for principal component analysis. `pathwayPCA` can analyze continuous, binary, and survival outcomes in studies with multiple covariate and/or interaction effects. We provide three case studies to illustrate pathway analysis with gene selection, integrative analysis of multi-omics datasets to identify driver genes, estimating and visualizing sample-specific pathway activities in ovarian cancer, and identifying sex-specific pathway effects in kidney cancer. `pathwayPCA` is an open source R package, freely available to the research community. We expect `pathwayPCA` to be a useful tool for empowering the wide scientific community on the analyses and interpretation of the wealth of multiomics data recently made available by TCGA, CPTAC and other large consortiums.

## INTRODUCTION

Pathway analysis has become a valuable strategy for analysing high-throughput -omics data. These pathway-based approaches test coordinated changes in functionally-related genes, which often belong to the same biological pathways. In addition to improving power by combining associated signals from multiple genes in the same pathway, these systems approaches can also shed more light on the underlying biological processes involved in diseases (1,2).

While many pathway analysis tools have been developed over the past decade, few of these tools can provide subject-specific estimates of pathway activities. However, to develop successful personalized treatment regimes, in addition to identifying disease-relevant pathways for the entire patient group, it is also important to identify individual patients with dysregulated pathway activities. Moreover, as technology advances, multiple types of -omics data across samples have become increasingly available. For example, the Cancer Genome Atlas (TCGA) and the Clinical Proteomic Tumor Analysis Consortium (CPTAC) have generated comprehensive molecular profiles including genomic, epigenomic, and proteomic expressions for human tumors (3,4). To perform integrative pathway analysis that leverages information in multi-omics datasets, measures of subject-specific pathway activity can be especially useful when examining relationships among multiple layers of cellular activities.

Principal Component Analysis (PCA) is a popular technique for reducing data dimensionality to capture variations in individual genes or subjects. In particular, principal components (PCs) have previously been used as sample-specific summaries of gene expression values from multiple genes (5). However, when the number of genes in the pathway is moderately large, genes unrelated to the phenotype may introduce noise and obscure the gene set association signal. Typically, only a subset of genes from an *a priori* defined pathway participate in the cellular process related to variations in phenotype, where each gene in the subset contributes a modest amount. Therefore, gene selection is an important issue in pathway analysis.

To address this challenge, we developed a supervised and an unsupervised approach for gene selection before performing PC-based pathway analysis (5-7). Both supervised PCA (SuperPCA) and Adaptive Elastic-net Sparse PCA (AES-PCA) perform gene selection to remove irrelevant genes before estimating pathway-specific PCs. In the SuperPCA approach (6,7), genes with an association measure below a certain threshold (estimated by cross-validation) are discarded, and the remaining genes are used to construct the PCs. To account for this gene selection process, a two-component mixture distribution based on the Gumbel Extreme Value distribution is used to estimate pathway *p*-values. In the unsupervised AES-PCA approach (5), PCs are estimated from a coherent subset of genes selected via the elastic-net penalty, which shrinks the estimates for many genes to zero (8).

Previously, both approaches have been shown to have superior performance when compared to popular pathway analysis approaches such as Hypergeometric test (9), GSEA (10), globalTest (11) in the analysis of gene expression data (7), GWAS data (6), and DNA methylation data (12). Furthermore, under the well-established PCA framework, both of these methodologies provide subject-specific estimates of pathway activities.

To make these powerful methodologies available to the wider research community, we present `pathwayPCA`, an R package that implements the SuperPCA and AES-PCA approaches for integrative pathway analysis. Table 1 includes a comparison of `pathwayPCA` with popular pathway analysis tools. The main features of `pathwayPCA` include:

**Table 1.**
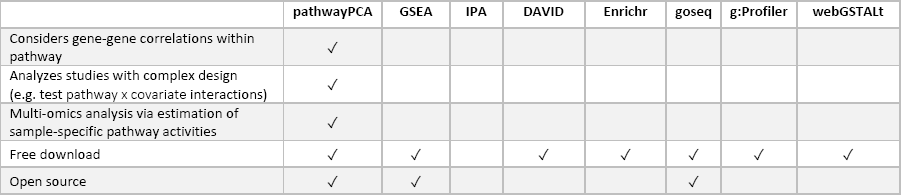
A comparison of different pathway analysis tools.

1. Performing pathway analysis for datasets with binary, continuous, or survival outcomes.
2. Extracting relevant genes from significant pathways using the SuperPCA and AES-PCA approaches.
3. Computing sample specific estimate of pathway activities. The PCs computed based on the selected genes are estimates of pathway activities for individual subjects. These estimated latent variables can then be used to perform integrative pathway analysis, such as multi-omics analysis.
4. Analyzing studies with complex experimental designs that include multiple covariates and/or interaction effects, e.g., testing if pathway associations with clinical phenotype are different between male and female subjects.
5. Performing analyses with enhanced computational efficiency via parallel computing and enhanced data safety via S4-class data objects.

## MATERIAL AND METHODS

`pathwayPCA` is licensed under GPL-3, is freely available to the general public, and is currently being submitted to the Bioconductor repository (13). Figure 1 shows a schematic overview of `pathwayPCA`. The software webpage (https://gabrielodom.github.io/pathwayPCA/) and Supplementary Text include in-depth tutorials on each step of the analyses, as well as visualization of analysis results. We describe the major functions in `pathwayPCA` next.

**Figure 1.**
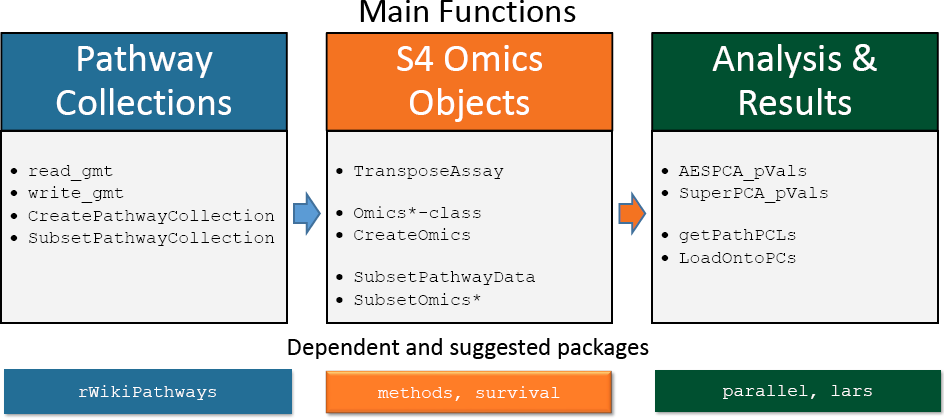
An overview of `pathwayPCA` functions.

### Creating data objects for pathway analysis

The `CreateOmics` function creates a S4 data object of class `Omics` based on several user input datasets: (1) an assay dataset, (2) a collection of pathways which can be imported by the `read_gmt` function, and (3) phenotype information for each sample, which can be binary, continuous, or survival outcome. Extensive data checking is performed to ensure valid data are imported. For example, the `CreateOmics` function checks for proper feature names, features with near-zero variance, overlap between features and the given pathway collection, and complete cases in the response. Because `pathwayPCA` implements a *self-contained test* (14), only genes in both assay data and pathway collection are considered for analysis. For each analysis, the `CreateOmics` function reports the number of genes in both assay data and pathway collection, as well as number of pathways with at least 5 or more genes (specified by parameter `minPathSize`).

### Testing pathway association with phenotype

Once we have a valid `Omics`-class object, we can perform pathway analysis using the AES-PCA or SuperPCA methods, which are implemented in the `AESPCA_pVals` and `SuperPCA_pVals` functions, respectively. Both functions return a table of the analyzed pathways sorted by p-values with additional fields including pathway name, description, number of included features, and estimated False Discovery Rate, as well as a list of the PCs and corresponding loadings for each pathway.

Briefly, in the AES-PCA method, we first extract latent variables (PCs) representing activities within each pathway using a dimension reduction approach based on adaptive, elastic-net, sparse PCA. The estimated latent variables are then tested against phenotypes using a linear regression model phenotype ∼ PC1 (default) or a permutation test that permutes sample labels. Note that the AES-PCA approach does not use response information to estimate pathway PCs, so it is an unsupervised approach.

On the other hand, SuperPCA is a supervised approach: the subset of genes most associated with disease outcome are used to estimate the latent variable for a pathway. Because of this gene selection step, the test statistic in SuperPCA model can no longer be approximated well using the Student’s *t*-distribution. To account for the gene selection step, `pathwayPCA` estimates *p*-values from a two-component mixture of Gumbel extreme value distributions instead (6,7).

### Extracting relevant genes and dataset for significant pathways

Because pathways are defined *a priori* (independently of the data), typically only a subset of genes within each pathway are relevant to the phenotype and contribute to a pathway’s significance. In our analyses, these relevant genes are the genes with nonzero loadings in the PCs extracted by AES-PCA or SuperPCA. Given results from the `AESPCA_pVals` and `SuperPCA_pVals` functions and a specific pathway name, the `getPathPCLs` function returns the loadings for each gene in the particular pathway. In addition, to allow for easy inspection of data and further in-depth analysis, the `SubsetPathwayData` function can be used to extract assay data for genes within a particular pathway, merged with the phenotype information.

### Estimating subject-specific pathway activities

In the study of complex diseases, there is often considerable heterogeneity among different subjects with regard to underlying causes of disease and benefit of particular treatment. Therefore, in addition to identifying disease-relevant pathways for the entire patient group, successful (personalized) treatment regimens will also depend upon knowing if a particular pathway is dysregulated for an individual patient. To this end, the `getPathPCLs` function can also extract sample estimates for the PCs, which allow users to assess pathway activities specifically for each patient. These subject-specific estimates for pathway activities can also facilitate multi-omics pathway analysis, as we will illustrate in Case Study 2 below.

## RESULTS

### *Case Study 1*: A WikiPathways analysis of ovarian cancer protein expression data

For this example, we downloaded a mass-spectrometry based global proteomics dataset generated by the Clinical Proteomic Tumor Analysis Consortium (CPTAC). The normalized protein abundance expression dataset for ovarian cancer was obtained from the LinkedOmics database at http://linkedomics.org/data_download/TCGA-OV/. We used the dataset “Proteome (PNNL, Gene level)” which was generated by the Pacific Northwest National Laboratory (PNNL). One subject was removed due to missing survival outcome. Missing protein expression values were imputed using the Bioconductor package `impute` under default settings (15). The final dataset consisted of 5162 protein expression values for 83 samples.

Using the `CreateOmics` function, we first grouped these protein expression values by pathways defined from the June 2018 WikiPathways (16) collection for *homo sapiens* (http://data.wikipathways.org/20180610/gmt/wikipathways-20180610-gmt-Homo_sapiens.gmt). The `AESPCA_pVals` function was then used to extract PC1 (the PC that accounts for most variation among samples) for each pathway. For each pathway, `AESPCA_pVals` fits overall survival using the Cox proportional hazards model with PC1 as predictor. Figure 2 shows the top 20 most significant pathways with *p*-values less than 0.0001.

**Figure 2.**
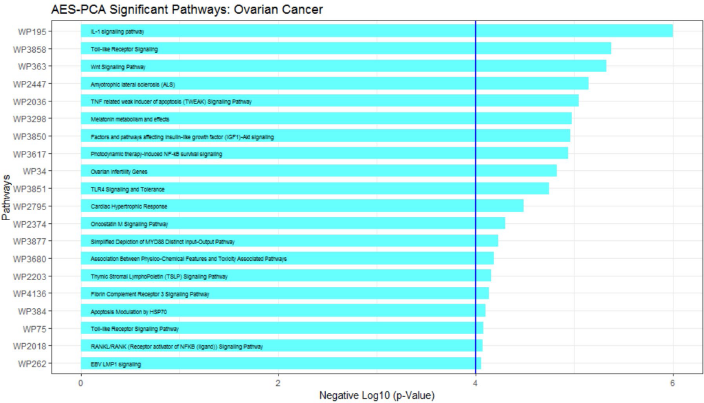
Most significant pathways by AES-PCA for ovarian cancer pathway analysis.

The most significant pathway is the IL-1 signaling pathway. To understand which proteins contributed most to pathway significance, the `getPathPCLs` function can extract the loadings for PC1 from this pathway (the weights of the proteins in the estimated PC1). Figure 3A provides a visualization for contributions of the relevant genes (IKBKB, NFKB1, MYD88) to PC1 in this pathway. In addition, the `getPathPCLs` function also returns subject-specific estimates of the first PC. Figure 3B shows the distribution of pathway activities in the IL-1 signaling pathway for all the subjects.

**Figure 3.**
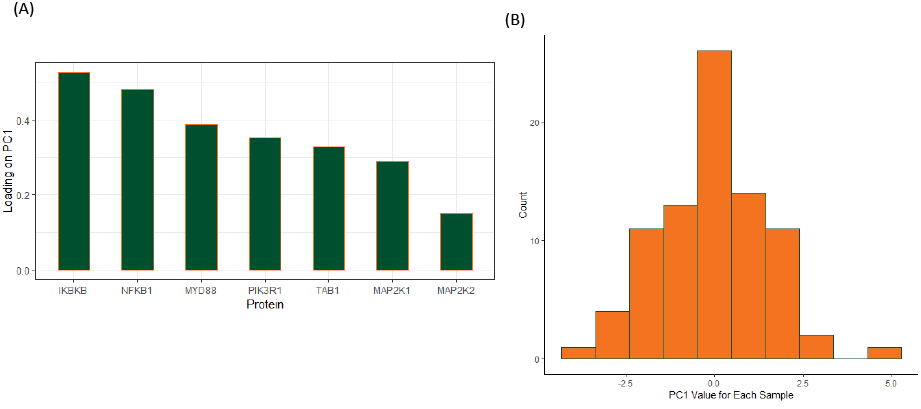
Gene specific and sample specific estimates for the `IL-1 signaling pathway`. (A) Relevant genes selected by AES-PCA. Shown are loadings of `PC1`, which are weights for each gene that contribute to `PC1` by AES-PCA. (B) Distribution of sample-specific estimate of pathway activities. Shown are estimated PC1 by AES-PCA for each sample.

Users are often also interested in examining the actual dataset used for analysis of the top pathways, especially for the relevant genes within the pathway. The `SubsetPathwayData` function extracts such a dataset with protein expressions and survival outcomes, matched by each sample for a given pathway. This pathway-specific dataset allows us to further explore the relevant genes in the pathway. For example, we can then fit a Cox regression model to individual genes (Figure 4A) or plot gene-specific Kaplan-Meier curves (Figure 4B).

**Figure 4.**
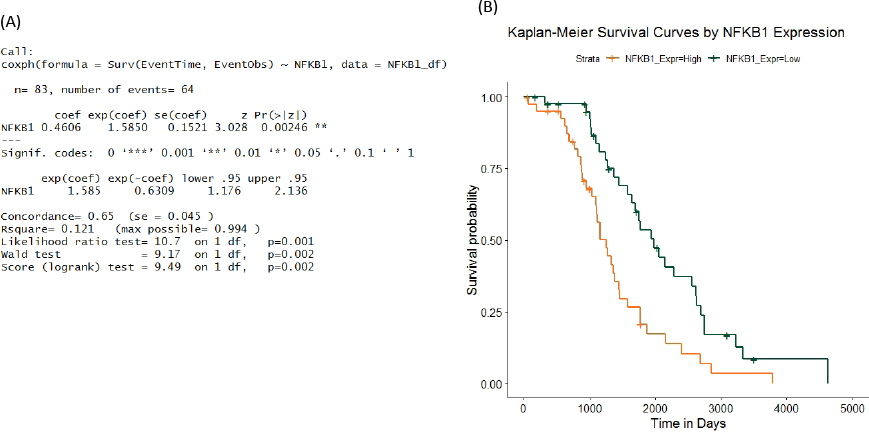
`pathwayPCA` provides functionality to extract data specific to a particular pathway or gene, which can be used for further in-depth analysis such as (A) Cox regression model (B) Kaplan-Meier survival curves.

Note that while we have illustrated an analysis using the AES-PCA methodology, the analysis workflow for the SuperPCA pathway analysis method is the same, except for replacing the `AESPCA_pVals` function call with a call to the `SuperPCA_pVals` function instead. In the results from these two different approaches, there will be slight discrepancies between the significant pathways identified and estimated loadings for individual proteins. This is because the gene-selection criteria used by the two methodologies are different. In AES-PCA, the focus is on groups of correlated genes, agnostic to phenotype; while in SuperPCA, the focus is on groups of genes most associated with phenotype. These two techniques in gene selection correspond to different biological hypotheses in how genes within a pathway influence outcomes. While the SuperPCA approach assumes the most significant genes within a pathway contribute most to the latent variable that captures pathway activity, users of the AES-PCA approach believe a coherent subset of genes, some of which might not be the most significant genes, contribute most to pathway activities.

### *Case Study 2*: An integrative multi-omics pathway analysis of ovarian cancer data

While copy number alterations are common genomic aberrations in ovarian cancer, recent studies have shown these changes do not necessarily lead to concordant changes in protein expression (17,18). In Case Study 1 above, we illustrated testing pathway activities in protein expression against survival outcome. In this section, we will additionally test pathway activities in copy number variations against survival outcome. Moreover, we will perform integrative analysis to identify those survival-associated protein pathways, genes, and samples likely driven by copy number alterations.

Briefly, the TCGA ovarian cancer gene-level copy number variation (CNV) data estimated using the GISTIC2 method (19) was downloaded from UCSC Xena Functional Genomics Browser (http://xena.ucsc.edu/) (20). The `CreateOmics` and `AESPCA_pvals` functions were used to identify CNV pathways significantly associated with survival outcomes. We identified 128 significant CNV pathways with *p*-values less than 0.05. A comparison with protein expression analysis showed 23 pathways were significantly associated with overall survival in both CNV and protein expressions data at 5% significance level (Figure 5).

**Figure 5.**
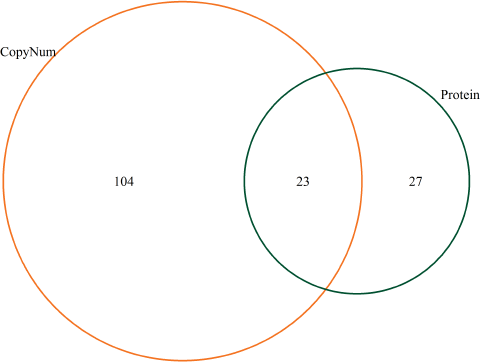
Pathways significantly associated (*p*-values < 0.05) with overall survival in copy number variations and protein expressions ovarian cancer datasets.

In addition, as illustrated in Case Study 1, the `getPathPCLs` function was used to identify genes and proteins with nonzero loadings in the IL-1 signaling pathway separately. These are the features that drive pathway significance in CNV and protein pathway analysis. The results showed that the NFKB1 gene had non-zero loadings and contributed to both CNV and protein pathway significance. A correlation plot (Figure 6) showed copy number variations for this gene were highly correlated with protein expression. This supported the hypothesis that the NFKB1 gene is likely a driver instead of passenger gene.

**Figure 6.**
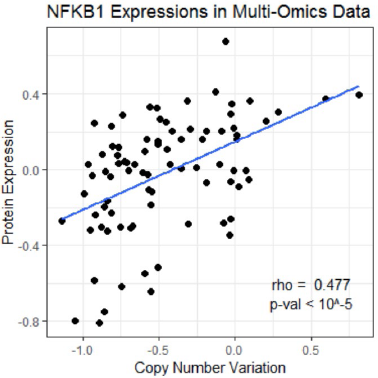
Significant association between copy number variations and protein expressions for the `NFKB1` gene, which were selected by AES-PCA in both copy number and protein expression pathway analysis of ovarian cancer data.

Figure 3B shows there can be considerable heterogeneity in pathway activities between the patients. One possible reason could be that copy number changes might not lead to changes in protein expression for some of the patients. The `getPathPCLs` function can be used to estimate pathway activities for each patient, for protein expressions and copy number expressions CNV separately. These estimates can then be visualized jointly using a Circos plot (21) constructed with the `circlize` package (22) (Figure 7).

**Figure 7.**
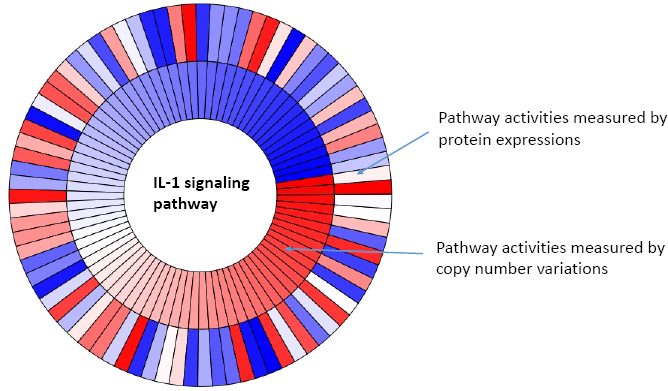
A Circos plot of normalized copy number (inner circle) and protein expression (outer circle) pathway activities for the `IL-1 signaling pathway` in the ovarian cancer dataset samples. Each bar correspond to a patient sample. Red color indicates higher expression values and more pathway activity for the sample. Blue color indicates lower expression values and lower pathway activity for the sample. Note that only some of the patients have concordant changes in copy number and protein expression.

### *Case Study 3*: An analysis of sex-specific pathway gene expression effects on kidney cancer

`pathwayPCA` is capable of analyzing complex studies with multiple experimental factors. In this case study, we will illustrate using `pathwayPCA` to test differential association of pathway activities with survival outcomes in male and female subjects. For many cancers, there are considerable sex disparities in the prevalence, prognosis, and treatment responses (23).

To understand the underlying biological differences that might contribute to the sex disparities in Cervical Kidney renal papillary cell carcinoma (KIRP), we downloaded the TCGA KIRP gene expression dataset from the Xena Functional Genomics browser (20) and tested sex × pathway activity interaction for each WikiPathway. Specifically, we organized the data using the `CreateOmics` function, estimated pathway activities for each subject using the `AESPCA_pVals` function, extracted the PCA results with the `getPathPCLs` function, and then fit the following Cox proportional hazards regression model to each pathway:

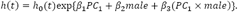

In this model, *h*(*t*) is expected hazard at time *t, h*_0_(*t*) is baseline hazard for the reference group, variable male is an indicator variable for male samples, and `PC1` is a pathway’s estimated first principal component based on AES-PCA. Table 2 shows there are 14 pathways with significant *p*-values less than 0.05 for the `PC1 × male` interaction, indicating the association of pathway gene expression (`PC1`) with survival for these pathways is highly dependent on sex of the subjects.

**Table 2.** Pathways with significant `Sex × Pathway` interaction in sex-specific pathway analysis of kidney cancer dataset.

| Terms | Description | Z | p-value |
| --- | --- | --- | --- |
| **WP1559** | TFs Regulate miRNAs related to cardiac hypertrophy | -2.76 | 0.00573 |
| **WP453** | Inflammatory Response Pathway | -2.5 | 0.0126 |
| **WP3893** | Development and heterogeneity of the ILC family | -2.4 | 0.0166 |
| **WP3929** | Chemokine signaling pathway | -2.32 | 0.0206 |
| **WP2849** | Hematopoietic Stem Cell Differentiation | -2.16 | 0.0305 |
| **WP1423** | Ganglio Sphingolipid Metabolism | -2.11 | 0.0346 |
| **WP3892** | Development of pulmonary dendritic cells and macrophage subsets | -2.1 | 0.036 |
| **WP3678** | Amplification and Expansion of Oncogenic Pathways as Metastatic Traits | -2.09 | 0.0363 |
| **WP4141** | PI3K/AKT/mTOR - VitD3 Signalling | 2.05 | 0.0407 |
| **WP3941** | Oxidative Damage | -2.01 | 0.0442 |
| **WP3863** | T-Cell antigen Receptor (TCR) pathway during Staphylococcus aureus infection | -1.99 | 0.0462 |
| **WP3872** | Regulation of Apoptosis by Parathyroid Hormone-related Protein | 1.98 | 0.0472 |
| **WP3672** | LncRNA-mediated mechanisms of therapeutic resistance | -1.98 | 0.0475 |
| **WP3967** | miR-509-3p alteration of YAP1/ECM axis | -1.98 | 0.0481 |

As an example, the pathway with the most significant `PC1 × male` interaction is the `TFs Regulate miRNAs related to cardiac hypertrophy` pathway. Cardiac hypertrophy, specifically let ventricular hypertrophy is highly prevalent in kidney disease patients (24). Gender differences have been observed in cardiac hypertrophy, which may be related to estrogens and testosterone (25). A recent integrative systems biology study showed that miRNA-mRNA network also plays an important role for gender differences in cardiac hypertrophy (26). The genes with large PC loadings in this identified pathway include `PPP3R1, STAT3 and TGFB1`, which regulate miRNA `hsa-mir-133b, hsa-mir-21` and `MIR29A`. In Figure 8, we grouped subjects by median `PC1` values for each sex. These Kaplan-Meier curves showed that while high or low pathway activities were not associated with survival in male subjects (green and purple curves, respectively), female subjects with high pathway activities (red) had significantly worse survival outcomes than those with low pathway activities (blue).

**Figure 8.**
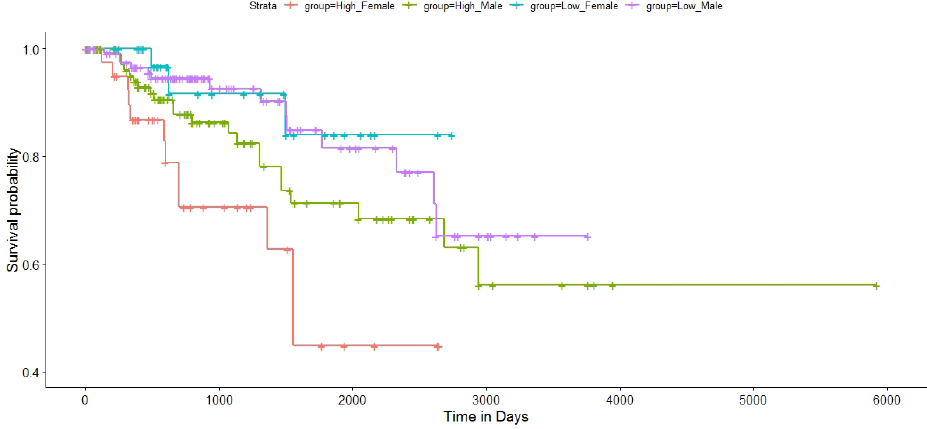
The Kaplan-Meier curves showed that while high or low pathway activities were not associated with survival in male subjects (green and purple curves, respectively), female subjects with high pathway activity (red) had significantly worse survival outcomes than those with low pathway activities (blue).

## DISCUSSION

Although pathway analysis has become a popular strategy for interpreting high-throughput -omics datasets, most of the available tools are limited to simple experimental designs and testing simple hypothesis that compares two groups. In particular, there is a lack of tools that can facilitate pathway analysis of multi-omics data, or provide sample-specific pathway activities. Here, we have presented `pathwayPCA`, a unique pathway analysis software that utilizes modern statistical methodology including supervised PCA and adaptive elastic-net PCA for principal component analysis and gene selection. The strength of `pathwayPCA` lies in its flexibility and versatility. It can be used to analyse studies with binary, continuous, or survival outcomes, as well as those with multiple covariate and/or interaction effects. Moreover, under the well-established PCA framework, contributions of individual genes toward pathway significance can be extracted and sample-specific pathway activities can be estimated. As we have illustrated in case studies, these functionalities can be especially helpful for visualizing pathway activities in multi-omics datasets and identifying driver genes. Computationally, `pathwayPCA` is efficient with options for parallel computing on all major operating systems. Testing 1000+ pathways typically takes only a few minutes.

## AVAILABILITY

`pathwayPCA` is freely available at GitHub at https://gabrielodom.github.io/pathwayPCA/ and is currently being submitted to the Bioconductor repository.

## REFERENCES

1. Garcia-Campos, M.A., Espinal-Enriquez, J. and Hernandez-Lemus, E. (2015) Pathway Analysis: State of the Art. Frontiers in physiology, 6, 383.

2. Wang, L., Jia, P., Wolfinger, R.D., Chen, X. and Zhao, Z. (2011) Gene set analysis of genomewide association studies: methodological issues and perspectives. Genomics, 98, 1–8.

3. Tomczak, K., Czerwinska, P. and Wiznerowicz, M. (2015) The Cancer Genome Atlas (TCGA): an immeasurable source of knowledge. Contemporary oncology, 19, A68–77.

4. Vasaikar, S.V., Straub, P., Wang, J. and Zhang, B. (2018) LinkedOmics: analyzing multi-omics data within and across 32 cancer types. Nucleic acids research, 46, D956–D963.

5. Chen, X. (2011) Adaptive elastic-net sparse principal component analysis for pathway association testing. Statistical applications in genetics and molecular biology, 10.

6. Chen, X., Wang, L., Hu, B., Guo, M., Barnard, J. and Zhu, X. (2010) Pathway-based analysis for genome-wide association studies using supervised principal components. Genetic epidemiology, 34, 716–724.

7. Chen, X., Wang, L., Smith, J.D. and Zhang, B. (2008) Supervised principal component analysis for gene set enrichment of microarray data with continuous or survival outcomes. Bioinformatics, 24, 2474–2481.

8. Zou, H. and Zhang, H.H. (2009) On the Adaptive Elastic-Net with a Diverging Number of Parameters. Annals of statistics, 37, 1733–1751.

9. Falcon, S. and Gentleman, R. (2007) Using GOstats to test gene lists for GO term association. Bioinformatics, 23, 257–258.

10. Subramanian, A., Tamayo, P., Mootha, V.K., Mukherjee, S., Ebert, B.L., Gillette, M.A., Paulovich, A., Pomeroy, S.L., Golub, T.R., Lander, E.S. et al. (2005) Gene set enrichment analysis: a knowledge-based approach for interpreting genome-wide expression profiles. Proceedings of the National Academy of Sciences of the United States of America, 102, 15545–15550.

11. Goeman, J.J., van de Geer, S.A., de Kort, F. and van Houwelingen, H.C. (2004) A global test for groups of genes: testing association with a clinical outcome. Bioinformatics, 20, 93–99.

12. Zhang, Q., Zhao, Y., Zhang, R., Wei, Y., Yi, H., Shao, F. and Chen, F. (2016) A Comparative Study of Five Association Tests Based on CpG Set for Epigenome-Wide Association Studies. PloS one, 11, e0156895.

13. Huber, W., Carey, V.J., Gentleman, R., Anders, S., Carlson, M., Carvalho, B.S., Bravo, H.C., Davis, S., Gatto, L., Girke, T. et al. (2015) Orchestrating high-throughput genomic analysis with Bioconductor. Nature methods, 12, 115–121.

14. Goeman, J.J. and Buhlmann, P. (2007) Analyzing gene expression data in terms of gene sets: methodological issues. Bioinformatics, 23, 980–987.

15. Hastie, T., Tibshirani, R., Narasimhan, B. and Chu, G. (2018), Bioconductor. 1.54.0 ed. Bioconductor.

16. Kutmon, M., Riutta, A., Nunes, N., Hanspers, K., Willighagen, E.L., Bohler, A., Melius, J., Waagmeester, A., Sinha, S.R., Miller, R. et al. (2016) WikiPathways: capturing the full diversity of pathway knowledge. Nucleic acids research, 44, D488–494.

17. Zhang, B., Wang, J., Wang, X., Zhu, J., Liu, Q., Shi, Z., Chambers, M.C., Zimmerman, L.J., Shaddox, K.F., Kim, S. et al. (2014) Proteogenomic characterization of human colon and rectal cancer. Nature, 513, 382–387.

18. Zhang, H., Liu, T., Zhang, Z., Payne, S.H., Zhang, B., McDermott, J.E., Zhou, J.Y., Petyuk, V.A., Chen, L., Ray, D. et al. (2016) Integrated Proteogenomic Characterization of Human High-Grade Serous Ovarian Cancer. Cell, 166, 755–765.

19. Mermel, C.H., Schumacher, S.E., Hill, B., Meyerson, M.L., Beroukhim, R. and Getz, G. (2011) GISTIC2.0 facilitates sensitive and confident localization of the targets of focal somatic copynumber alteration in human cancers. Genome biology, 12, R41.

20. Goldman, M., Craft, B., Kamath, A., Brooks, A.N., Zhu, J. and Haussler, D. (2018) The UCSC Xena Platform for cancer genomics data visualization and interpretation. bioRxiv.

21. Krzywinski, M., Schein, J., Birol, I., Connors, J., Gascoyne, R., Horsman, D., Jones, S.J. and Marra, M.A. (2009) Circos: an information aesthetic for comparative genomics. Genome Res, 19, 1639–1645.

22. Gu, Z., Gu, L., Eils, R., Schlesner, M. and Brors, B. (2014) circlize Implements and enhances circular visualization in R. Bioinformatics, 30, 2811–2812.

23. Yuan, Y., Liu, L., Chen, H., Wang, Y., Xu, Y., Mao, H., Li, J., Mills, G.B., Shu, Y., Li, L. et al. (2016) Comprehensive Characterization of Molecular Differences in Cancer between Male and Female Patients. Cancer cell, 29, 711–722.

24. Taddei, S., Nami, R., Bruno, R.M., Quatrini, I. and Nuti, R. (2011) Hypertension, left ventricular hypertrophy and chronic kidney disease. Heart failure reviews, 16, 615–620.

25. Regitz-Zagrosek, V., Oertelt-Prigione, S., Seeland, U. and Hetzer, R. (2010) Sex and gender differences in myocardial hypertrophy and heart failure. Circulation journal : official journal of the Japanese Circulation Society, 74, 1265–1273.

26. Harrington, J., Fillmore, N., Gao, S., Yang, Y., Zhang, X., Liu, P., Stoehr, A., Chen, Y., Springer, D., Zhu, J. et al. (2017) A Systems Biology Approach to Investigating Sex Differences in Cardiac Hypertrophy. Journal of the American Heart Association, 6.

