## Supplementary file for "pathwayPCA: an R package for integrative pathway analysis with modern PCA methodology and gene selection"

### Integrative Pathway Analysis with pathwayPCA

2019-02-06

#### 1. Introduction

`pathwayPCA` is an integrative analysis tool that implements the principal component analysis (PCA) based pathway analysis approaches described in Chen et al. (2008), Chen et al. (2010), and Chen (2011). `pathwayPCA` allows users to:

1. Test pathway association with binary, continuous, or survival phenotypes.
2. Extract relevant genes in the pathways using the SuperPCA and AESPCA approaches.
3. Compute *principal components* (PCs) based on the selected genes. These estimated latent variables represent pathway activities for individual subjects, which can then be used to perform integrative pathway analysis, such as multi-omics analysis.

4. Extract relevant genes that drive pathway significance as well as data corresponding to these relevant genes for additional in-depth analysis.
5. Perform analyses with enhanced computational efficiency with parallel computing and enhanced data safety with S4-class data objects.
6. Analyze studies with complex experimental designs, with multiple covariates, and with interaction effects, e.g., testing whether pathway association with clinical phenotype is different between male and female subjects.

The development version of the pathwayPCA package can be installed from GitHub:

```
devtools::install_github("gabrielodom/pathwayPCA")
```

Throughout this vignette, we will make use of the `tidyverse` suite of utility packages (https://www.tidyverse.org/). The `tidyverse` and `pathwayPCA` can be loaded into R using:

```
library(tidyverse)
library(pathwayPCA)
```

### 2. Case study: identifying significant pathways in protein expressions associated with survival outcome in ovarian cancer data

#### 2.1. Ovarian cancer dataset

For this example, we will use the mass spectrometry based global proteomics data for ovarian cancer recently generated by the Clinical Proteomic Tumor Analysis Consortium (CPTAC). The normalized protein abundance expression dataset can be obtained from the LinkedOmics database at http://linkedomics.org/data_download/TCGA-OV/. We used the dataset "Proteome (PNNL, Gene level)" which was generated by the Pacific Northwest National Laboratory (PNNL). One subject was removed due to missing survival outcome. Missing values in proteins expression data were imputed using the Bioconductor package `impute` under default settings. The final dataset consisted of 5162 protein expression values for 83 samples.

#### 2.2. Creating an `Omics` data object for pathway analysis

First, we need to create an `Omics`-class data object that stores

(1) the expression dataset
(2) phenotype information for the samples
(3) a collection of pathways

##### 2.2.1 Expression and Phenotype Data

We can obtain datasets 1 and 2 for the ovarian cancer dataset by loading the `ovarian_PNNL_survival.RDS` data file included in the `pathwayPCA` package.

```
dataDir_path <- system.file(
  "extdata", package = "pathwayPCA", mustWork = TRUE
)
ovSurv_df <- readRDS(paste0(dataDir_path, "/ovarian_PNNL_survival.RDS"))
```

The `ovSurv_df` dataset is a data frame with protein expression levels and survival outcome matched by sample IDs. The variables (columns) include overall survival time and censoring status, as well as expression data for 5162 proteins for each of the 83 samples.

```
ovSurv_df [1:5, 1:5]
#> # A tibble: 5 x 5
#>   Sample       OS_time OS_death    A1BG      A2M
#>   <chr>          <int>    <int>   <dbl>    <dbl>
#> 1 TCGA.09.1664    2279        1  0.336  -0.00505
#> 2 TCGA.13.1484    3785        1 -0.848  -0.434
#> 3 TCGA.13.1488    2154        1 -0.0730  0.172
#> 4 TCGA.13.1489    2553        1  0.0154 -0.419
#> 5 TCGA.13.1494    2856        1 -0.495   0.112
```

##### 2.2.2 Pathway Collections

For the collection of pathways in (3), we need to specify a `.gmt` file, a text file with each row corresponding to one pathway. Each row contains an ID (column `TERMS`), an optional description (column `description`), and the genes in the pathway (all subsequent columns). Pathway collections in `.gmt` form can be downloaded from the MSigDB database at http://software.broadinstitute.org/gsea/msigdb/collections.jsp.

For WikiPathways, one can download monthly data releases in `.gmt` format using the `dowloadPathwayArchive()` function in the `rWikiPathways` package from Bioconductor. For example, the following commands downloads the latest release of the human pathways from WikiPathways to your current directory:

```
library(rWikiPathways)
# library(XML) # necessary if you encounter an error with readHTMLTable
downloadPathwayArchive(
  organism = "Homo sapiens", format = "gmt"
)

trying URL 'http://data.wikipathways.org/current/gmt/wikipathways-20190110-gm
t-Homo_sapiens.gmt'
Content type '' length 174868 bytes (170 KB)
downloaded 170 KB

#> [1] "wikipathways-20190110-gmt-Homo_sapiens.gmt"
```

`pathwayPCA` includes the June 2018 Wikipathways collection for *homo sapiens*, which can be loaded using the `read_gmt` function:

```
wikipathways_PC <- read_gmt(
  paste0(dataDir_path, "/wikipathways_human_symbol.gmt"),
```

```
  description = TRUE
)
```

##### 2.2.3 Create an `OmicsSurv` Data Container

Now that we have these three data components, we create an `OmicsSurv` data container. Note that when `assayData_df` and `response` are supplied from two different files, the user must match and merge these data sets by sample IDs.

```
ov_OmicsSurv <- CreateOmics(
  # protein expression data
  assayData_df = ovSurv_df[, -(2:3)],
  # pathway collection
  pathwayCollection_ls = wikipathways_PC,
  # survival phenotypes
  response = ovSurv_df[, 1:3],
  # phenotype is survival data
  respType = "survival",
  # retain pathways with > 5 proteins
  minPathSize = 5
)

  ======  Creating object of class OmicsSurv  =======
The input pathway database included 5831 unique features.
The input assay dataset included 5162 features.
Only pathways with at least 5 or more features included in the assay dataset are
  tested (specified by minPathSize parameter). There are 324 pathways which meet
  this criterion.
Because pathwayPCA is a self-contained test (PMID: 17303618), only features in
  both assay data and pathway database are considered for analysis. There are 2064
  such features shared by the input assay and pathway database.
```

To see a summary of the `Omics` data object we just created, simply type the name of the object:

```
ov_OmicsSurv
#> Formal class 'OmicsSurv' [package "pathwayPCA"] with 6 slots
#>   ..@ eventTime           : num [1:83] 2279 3785 2154 2553 2856 ...
#>   ..@ eventObserved       : logi [1:83] TRUE TRUE TRUE TRUE TRUE TRUE ...
#>   ..@ assayData_df        :Classes 'tbl_df', 'tbl' and 'data.frame':    83 obs. of  5162 variables:
#>   ..@ sampleIDs_char      : chr [1:83] "TCGA.09.1664" "TCGA.13.1484" "TCGA.13.1488" "TCGA.13.1489" ...
#>   ..@ pathwayCollection   :List of 4
#>   .. ..- attr(*, "class")= chr [1:2] "pathwayCollection" "list"
```

```
#>   ..@ trimPathwayCollection:List of 5
#>   .. ..- attr(*, "class")= chr [1:2] "pathwayCollection" "list"
```

#### 2.3. Testing pathway association with phenotypes

##### 2.3.1 Method Description

Once we have a valid `Omics` object, we can perform pathway analysis using the AES-PCA (Adaptive, Elastic-net, Sparse PCA) or Supervised PCA (SuperPCA) methodology described in Chen et al. (2008), Chen et al. (2010), and Chen (2011).

Briefly, in the AES-PCA method, we first extract PCs representing activities within each pathway using a dimension reduction approach based on adaptive, elastic-net, sparse principal component analysis (https://doi.org/10.2202/1544-6115.1697). The estimated latent variables are then tested against phenotypes using a permutation test that permutes sample labels. Note that the AESPCA approach does not use the response information to estimate pathway PCs, so it is an unsupervised approach.

This is in contrast to the SuperPCA approach, where a selected subset of genes most associated with disease outcome are used to estimate the latent variable for a pathway (https://doi.org/10.1002/gepi.20532). Because of this gene selection step, the test statistics from the SuperPCA model can no longer be approximated well using the Student's $t$-distribution. To account for the gene selection step, `pathwayPCA` estimates $p$-values from a two-component mixture of Gumbel extreme value distributions instead.

##### 2.3.2 Implementation

Because the syntax for performing SuperPCA is nearly identical to the AES-PCA syntax, we will illustrate only the AES-PCA workflow below.

Note that when the value supplied to the `numReps` argument is greater than 0, the `AESPCA_pvals()` function employs a parametric test when estimating pathway significance via the following model: "phenotype ~ intercept + PC1". Pathway $p$-values are estimated based on a likelihood ratio test that compares this model to a null model (with intercept only).

```
ovarian_aespcOut <- AESPCA_pVals(
  # The Omics data container
  object = ov_OmicsSurv,
  # One principal component per pathway
  numPCs = 1,
  # Use parallel computing with 2 cores
  parallel = TRUE,
  numCores = 2,
  # # Use serial computing
  # parallel = FALSE,
  # Estimate the p-values parametrically
  numReps = 0,
  # Control FDR via Benjamini-Hochberg
```

```
  adjustment = "BH"
)
Part 1: Calculate Pathway AES-PCs
Initializing Computing Cluster: DONE
Extracting Pathway PCs in Parallel: DONE

Part 2: Calculate Pathway p-Values
Initializing Computing Cluster: DONE
Extracting Pathway p-Values in Parallel: DONE

Part 3: Adjusting p-Values and Sorting Pathway p-Value Data Frame
DONE
```

This ovarian_aespcOut object contains 3 components: a table of pathway $p$-values, AESPCA-estimated PCs of each sample from each pathway, and the loadings of each protein onto the AESPCs.

```
names(ovarian_aespcOut)
#> [1] "pVals_df"    "PCs_ls"      "loadings_ls"
```

##### 2.3.3 The pathway analysis results

For this ovarian cancer dataset, the top 10 most significant pathways identified by AES-PCA are:

```
ovarian_aespcOut$pVals_df[, -1]
#> # A tibble: 324 x 5
#>    n_tested terms  description                                 rawp FDR_BH
#>       <dbl> <chr>  <chr>                                      <dbl>  <dbl>
#>  1       30 WP195  IL-1 signaling pathway                   0.00247  0.279
#>  2       16 WP3858 Toll-like Receptor Signaling             0.00463  0.279
#>  3       18 WP363  Wnt Signaling Pathway                    0.00485  0.279
#>  4       24 WP2447 Amyotrophic lateral sclerosis (ALS)      0.00582  0.279
#>  5       22 WP2036 TNF related weak inducer of apoptosis (T~ 0.00641  0.279
#>  6       12 WP3298 Melatonin metabolism and effects         0.00688  0.279
#>  7       13 WP3850 Factors and pathways affecting insulin-l~ 0.00700  0.279
#>  8       10 WP3617 Photodynamic therapy-induced NF-kB survi~ 0.00714  0.279
#>  9        8 WP34   Ovarian Infertility Genes                0.00801  0.279
#> 10       14 WP3851 TLR4 Signaling and Tolerance             0.00870  0.279
#> # ... with 314 more rows
```

In this table, n_tested indicates the number of genes in both the pathway and assay data, i.e. the number of genes in a pathway that we actually tested for association with phenotype in AESPCA.

Before constructing a graph of the $p$-values, we modify the data frame of pathway $p$-values using the following commands:

```
ovOutGather_df <-
  # Take in the results data frame,
  ovarian_aespcOut$pVals_df %>%
```

```
  # add the score variable,
  mutate(score = -log(rawp)) %>%
  # subset the top 20 pathways, and
  slice(1:20) %>%
  # remove the columns we don't need
  select(-pathways, -n_tested, -rawp, -FDR_BH)
```

Now we plot the pathway significance level for the top 20 pathways.

```
ggplot(ovOutGather_df) +
  # set overall appearance of the plot
  theme_bw() +
  # Define the dependent and independent variables
  aes(x = reorder(terms, score), y = score) +
  # From the defined variables, create a vertical bar chart
  geom_col(position = "dodge", fill = "#66FFFF", width = 0.7) +
  # Add pathway labels
  geom_text(
    aes(x = reorder(terms, score), label = reorder(description, score), y = 0
.1),
    color = "black",
    size = 2,
    hjust = 0
  ) +
  # Set main and axis titles
  ggtitle("AES-PCA Significant Pathways: Ovarian Cancer") +
  xlab("Pathways") +
  ylab("Negative Log10 (p-Value)") +
  # Add a line showing the alpha = 0.0001 level
  geom_hline(yintercept = -log10(0.0001), size = 1, color = "blue") +
  # Flip the x and y axes
  coord_flip()
```

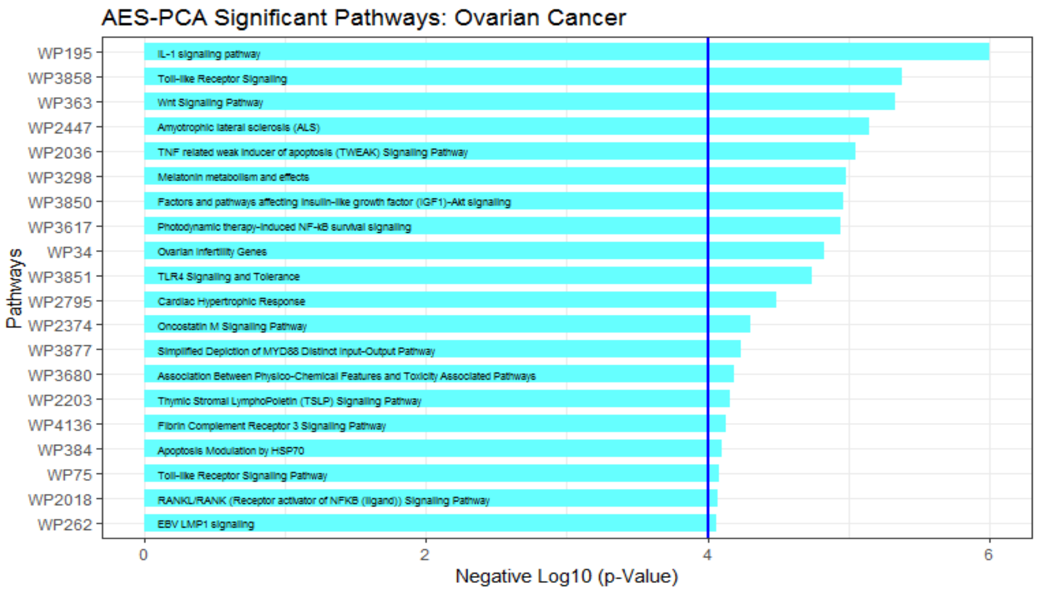

##### 2.3.4 Extract relevant genes from significant pathways

Because pathways are defined *a priori*, typically only a subset of genes within each pathway are relevant to the phenotype and contribute to pathway significance. In AESPCA, these relevant genes are the genes with nonzero loadings in the first PC of AESPCs.

For example, for the "IL-1 signaling pathway" (Wikipathways WP195), we can extract the **PC**s and their protein **L**oadings using the `getPathPCLs()` function:

```
wp195PCLs_ls <- getPathPCLs(ovarian_aespcOut, "WP195")
wp195PCLs_ls
#> $PCs
#> # A tibble: 83 x 2
#>    sampleID         V1
#>    <chr>         <dbl>
#>  1 TCGA.09.1664 -2.66
#>  2 TCGA.13.1484 -0.459
#>  3 TCGA.13.1488  0.603
#>  4 TCGA.13.1489 -1.70
#>  5 TCGA.13.1494 -0.726
#>  6 TCGA.13.1495 -0.316
#>  7 TCGA.13.1499  0.252
#>  8 TCGA.23.1123 -0.679
#>  9 TCGA.23.1124  1.83
#> 10 TCGA.24.1103 -1.93
#> # ... with 73 more rows
#>
#> $Loadings
#> # A tibble: 30 x 2
#>    featureID   PC1
#>    <chr>     <dbl>
#>  1 ATF2      0
#>  2 MAPK14    0
#>  3 AKT1      0
#>  4 IKBKB     0.527
#>  5 NFKB1     0.483
#>  6 PIK3R1    0.352
#>  7 PLCG1     0
#>  8 MAPK1     0
#>  9 MAPK9     0
#> 10 MAP2K1    0.289
#> # ... with 20 more rows
#>
#> $pathway
#> [1] "path22"
#>
#> $term
#> [1] "WP195"
#>
```

```
#> $description
#> [1] "IL-1 signaling pathway"
```

The proteins with non-zero loadings can be extracted using the following lines:

```
wp195Loadings_df <-
  wp195PCLs_ls$Loadings %>%
  filter(PC1 != 0)
wp195Loadings_df
#> # A tibble: 7 x 2
#>   featureID   PC1
#>   <chr>      <dbl>
#> 1 IKBKB      0.527
#> 2 NFKB1      0.483
#> 3 PIK3R1     0.352
#> 4 MAP2K1     0.289
#> 5 MAP2K2     0.150
#> 6 TAB1       0.328
#> 7 MYD88      0.389
```

We can also prepare these loadings for graphics:

```
wp195Loadings_df <-
  wp195Loadings_df %>%
  # Sort Loading from Strongest to Weakest
  arrange(desc(abs(PC1))) %>%
  # Order the Genes by Loading Strength
  mutate(featureID = factor(featureID, levels = featureID)) %>%
  # Add Directional Indicator (for Colour)
  mutate(Direction = factor(ifelse(PC1 > 0, "Up", "Down")))
```

Now we will construct a **col**umn chart with ggplot2's `geom_col()` function.

```
ggplot(data = wp195Loadings_df) +
  # Set overall appearance
  theme_bw() +
  # Define the dependent and independent variables
  aes(x = featureID, y = PC1, fill = Direction) +
  # From the defined variables, create a vertical bar chart
  geom_col(width = 0.5, fill = "#005030", color = "#f47321") +
  # Set main and axis titles
  labs(
    title = "Gene Loadings on IL-1 Signaling Pathway",
    x = "Protein",
    y = "Loading on PC1"
  ) +
  # Remove the legend
  guides(fill = FALSE)
```

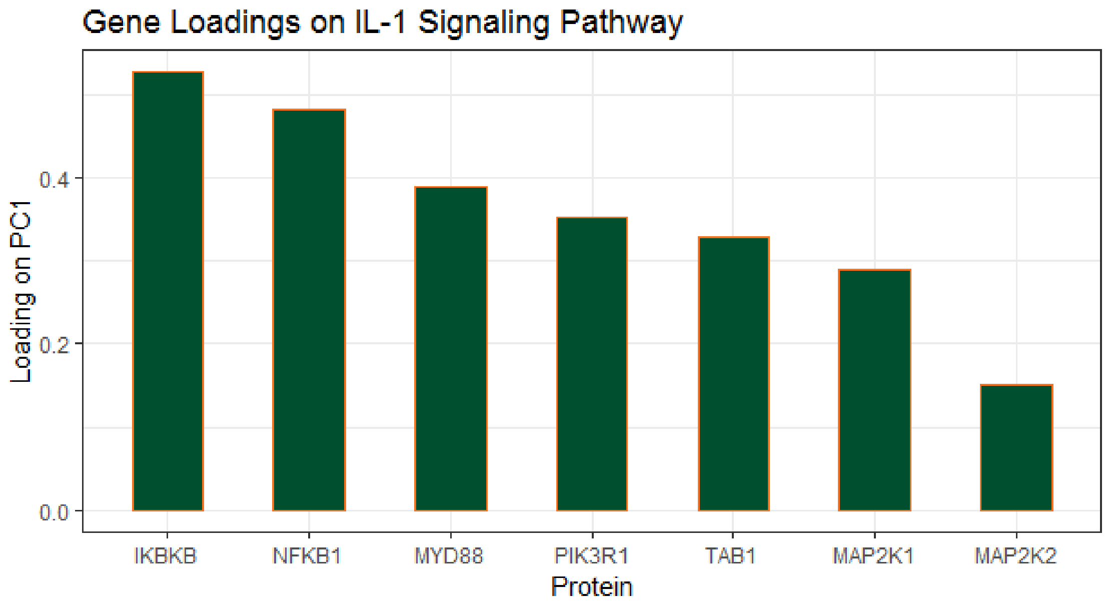

Alternatively, we can also plot the correlation of each gene with first PC for each gene. These correlations can be computed by using the `TidyCorrelation()` function in Section 3.3 of additional vignette "Chapter 5 - Visualizing the Results".

##### 2.3.5 Subject-specific PCA estimates

In the study of complex diseases, there is often considerable heterogeneity among different subjects with regard to underlying causes of disease and benefit of particular treatment. Therefore, in addition to identifying disease-relevant pathways for the entire patient group, successful (personalized) treatment regimens will also depend upon knowing if a particular pathway is dysregulated for an individual patient.

To this end, we can also assess subject-specific pathway activity. As we saw earlier, the `getPathPCLs()` function also returns subject-specific estimates for the individual pathway PCs.

```
ggplot(data = wp195PCLs_ls$PCs) +
  # Set overall appearance
  theme_classic() +
  # Define the independent variable
  aes(x = V1) +
  # Add the histogram layer
  geom_histogram(bins = 10, color = "#005030", fill = "#f47321") +
  # Set main and axis titles
  labs(
    title = "Distribution of Sample-specific Estimate of Pathway Activities",
    subtitle = paste0(wp195PCLs_ls$pathway, ": ", wp195PCLs_ls$description),
    x = "PC1 Value for Each Sample",
    y = "Count"
  )
```

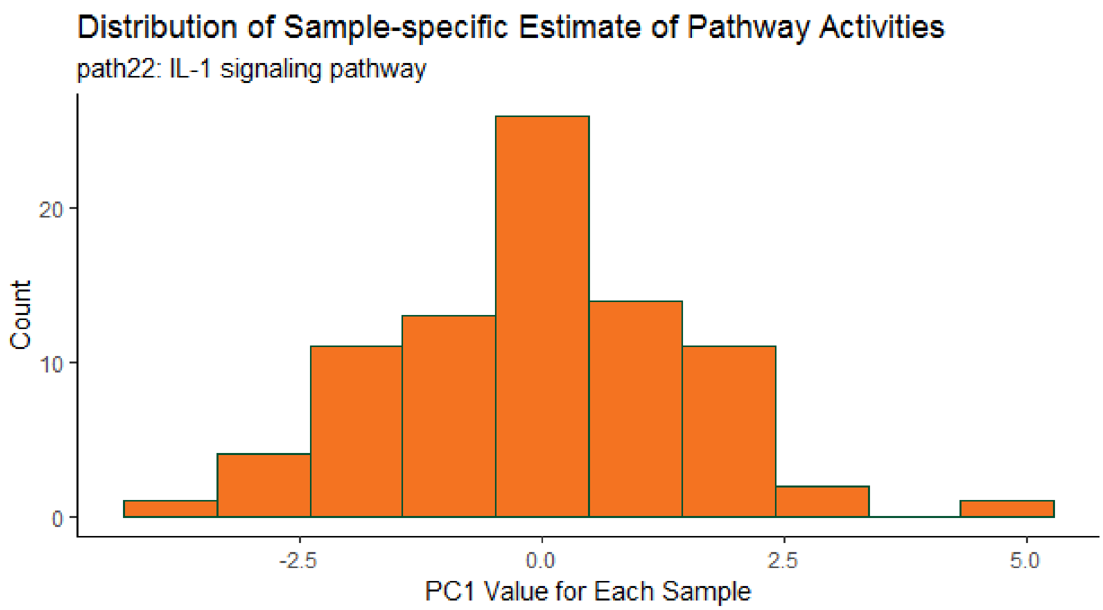

This graph shows there can be considerable heterogeneity in pathway activities between the patients.

##### 2.3.6 Extract analysis dataset for significant pathways

Users are often also interested in examining the actual data used for analysis of the top pathways, especially for the relevant genes with the pathway. To extract this dataset, we can use the `SubsetPathwayData()` function. These commands extract data for the most significant pathway (IL-1 signaling):

```
wp195Data_df <- SubsetPathwayData(ov_OmicsSurv, "WP195")
wp195Data_df
#> # A tibble: 83 x 33
#>    sampleID EventTime EventObs   ATF2  MAPK14    AKT1   IKBKB   NFKB1
#>    <chr>        <dbl> <lgl>     <dbl>   <dbl>   <dbl>   <dbl>   <dbl>
#>  1 TCGA.09~      2279 TRUE      0.623 -0.995   0.958  -1.71   -0.730
#>  2 TCGA.13~      3785 TRUE      1.17   1.71    0.575  -0.751   0.783
#>  3 TCGA.13~      2154 TRUE     -0.130 -0.127  -0.542  -0.915   0.418
#>  4 TCGA.13~      2553 TRUE      0.482 -0.0588 -0.292  -0.422  -0.488
#>  5 TCGA.13~      2856 TRUE      0.130  0.610   1.25   -0.950   0.393
#>  6 TCGA.13~      2749 TRUE     -0.823  0.266   1.79   -0.522  -0.0498
#>  7 TCGA.13~      3500 FALSE    -0.704  0.516  -0.519  -0.494   0.104
#>  8 TCGA.23~      1018 TRUE      0.270 -0.419  -3.63    0.0331 -0.114
#>  9 TCGA.23~      1768 TRUE     -2.61  -1.81   -0.0214  0.811   1.17
#> 10 TCGA.24~      1646 TRUE      0.270 -0.622   1.23   -0.924  -0.954
#> # ... with 73 more rows, and 25 more variables: PIK3R1 <dbl>, PLCG1 <dbl>,
#> #   MAPK1 <dbl>, MAPK9 <dbl>, MAP2K1 <dbl>, MAP2K2 <dbl>, MAP2K6 <dbl>,
#> #   PTPN11 <dbl>, REL <dbl>, RELA <dbl>, MAP3K7 <dbl>, IKBKG <dbl>,
#> #   TAB1 <dbl>, IRAK1 <dbl>, MYD88 <dbl>, IRAK4 <dbl>, ECSIT <dbl>,
#> #   TOLLIP <dbl>, MAPK3 <dbl>, MAP2K3 <dbl>, MAP2K4 <dbl>, TRAF6 <dbl>,
#> #   UBE2N <dbl>, SQSTM1 <dbl>, MAPKAPK2 <dbl>
```

We can also perform analysis for individual genes belonging to the pathway:

```
library(survival)
NFKB1_df <-
  wp195Data_df %>%
  select(EventTime, EventObs, NFKB1)

wp195_mod <- coxph(
  Surv(EventTime, EventObs) ~ NFKB1,
  data = NFKB1_df
)

summary(wp195_mod)
#> Call:
#> coxph(formula = Surv(EventTime, EventObs) ~ NFKB1, data = NFKB1_df)
#>
#>   n= 83, number of events= 64
#>
#>         coef exp(coef) se(coef)     z Pr(>|z|)
#> NFKB1 0.4606    1.5850   0.1521 3.028  0.00246 **
#> ---
#> Signif. codes:  0 '***' 0.001 '**' 0.01 '*' 0.05 '.' 0.1 ' ' 1
#>
#>       exp(coef) exp(-coef) lower .95 upper .95
#> NFKB1     1.585     0.6309     1.176     2.136
#>
#> Concordance= 0.65  (se = 0.045 )
#> Rsquare= 0.121   (max possible= 0.994 )
#> Likelihood ratio test= 10.7  on 1 df,   p=0.001
#> Wald test            = 9.17  on 1 df,   p=0.002
#> Score (logrank) test = 9.49  on 1 df,   p=0.002
```

Additionally, we can estimate Kaplan-Meier survival curves for patients with high or low expression values for individual genes:

```
# Add the direction
NFKB1_df <-
  NFKB1_df %>%
  mutate(NFKB1_Expr = ifelse(NFKB1 > median(NFKB1), "High", "Low"))

# Fit the survival model
NFKB1_fit <- survfit(
  Surv(EventTime, EventObs) ~ NFKB1_Expr,
  data = NFKB1_df
)
```

Finally, we can plot these K-M curves over NFKB1 protein expression.

```
library(survminer)
```

```
ggsurvplot(
  NFKB1_fit,
  conf.int = FALSE, pval = TRUE,
  xlab = "Time in Days",
  palette = c("#f47321", "#005030")
)
```

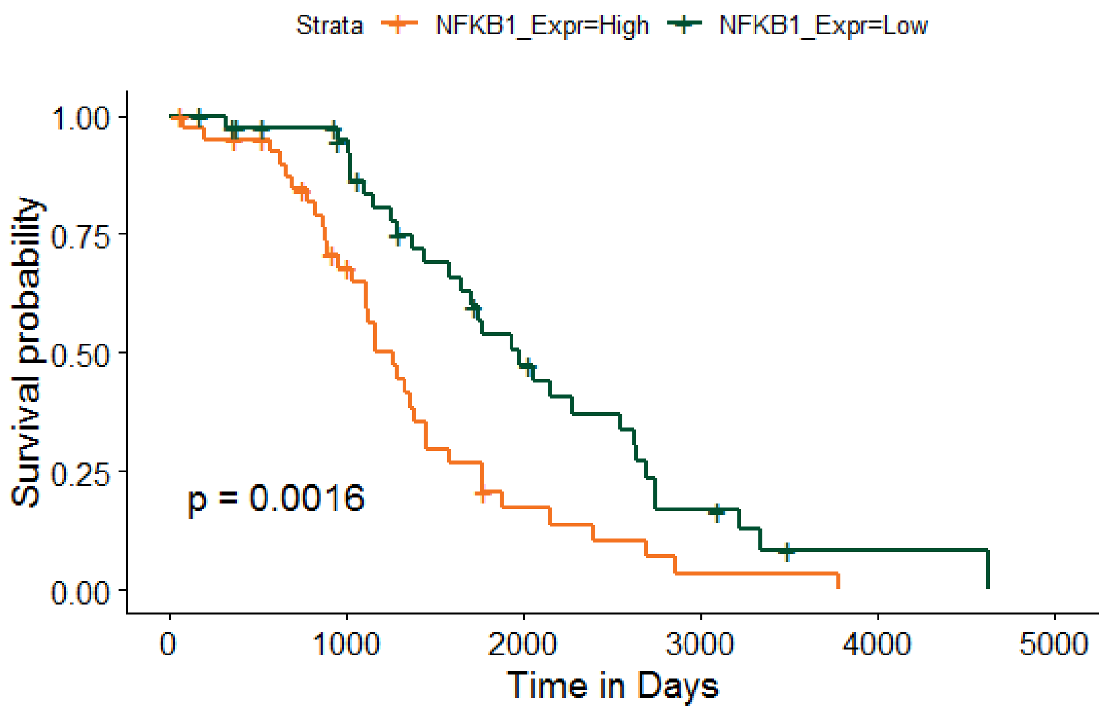

#### 3. Case study: an integrative multi-omics pathway analysis of ovarian cancer data

While copy number alterations are common genomic aberrations in ovarian carcer, recent studies have shown these changes do not necessarily lead to concordant changes in protein expression. In Section 2.3 above, we illustrated testing pathway activities in protein expression against survival outcome. In this section, we will additionally test pathway activities in copy number against survival outcome. Moreover, we will perform integrative analysis to identify those survival -associated protein pathways, genes, and samples driven by copy number alterations.

##### 3.1 Creating copy number `Omics` data object for pathway analysis

We can identify copy number (CNV) pathways significantly associated with survival in the same way as we did for protein expressions. This gene level CNV data was downloaded from UCSC Xena Functional Genomics Browser (http://xena.ucsc.edu/).

```
copyNumberClean_df <- readRDS(paste0(dataDir_path, "/OV_surv_x_CNV2.RDS"))
```

And now we create an `Omics` data container.

```
ovCNV_Surv <- CreateOmics(
  assayData_df = copyNumberClean_df[, -(2:3)],
  pathwayCollection_ls = wikipathways_PC,
  response = copyNumberClean_df[, 1:3],
  respType = "survival",
  minPathSize = 5
)

  ======  Creating object of class OmicsSurv  =======
The input pathway database included 5831 unique features.
The input assay dataset included 24776 features.
Only pathways with at least 5 or more features included in the assay dataset are
  tested (specified by minPathSize parameter). There are 424 pathways which meet
  this criterion.
Because pathwayPCA is a self-contained test (PMID: 17303618), only features in
  both assay data and pathway database are considered for analysis. There are
5637 such features shared by the input assay and pathway database.
```

Finally, we can apply the AESPCA method to this copy-number data container. Due to the large sample size, this will take a few moments.

```
ovCNV_aespcOut <- AESPCA_pVals(
  object = ovCNV_Surv,
  numPCs = 1,
  parallel = TRUE,
  numCores = 20,
  numReps = 0,
  adjustment = "BH"
)

Part 1: Calculate Pathway AES-PCs
Initializing Computing Cluster: DONE
Extracting Pathway PCs in Parallel: DONE

Part 2: Calculate Pathway p-Values
Initializing Computing Cluster: DONE
Extracting Pathway p-Values in Parallel: DONE

Part 3: Adjusting p-Values and Sorting Pathway p-Value Data Frame
DONE
```

#### 3.2 Identifying significant pathways and relevant genes in both CNV and protein level

Next, we identify the intersection of significant pathways based on both CNV and protein data. First, we will create a data frame of the pathway $p$-values from both CNV and proteomics.

```
# Copy Number
CNVpvals_df <-
  ovCNV_aespcOut$pVals_df %>%
  mutate(rawp_CNV = rawp) %>%
  select(description, rawp_CNV) %>%
  filter(rawp_CNV < 0.05)

# Proteomics
PROTpvals_df <-
  ovarian_aespcOut$pVals_df %>%
  mutate(rawp_PROT = rawp) %>%
  select(terms, description, rawp_PROT) %>%
  filter(rawp_PROT < 0.05)

# Intersection
SigBoth_df <- inner_join(PROTpvals_df, CNVpvals_df, by = "description")
# WnT Signaling Pathway is listed as WP363 and WP428
```

The results showed there are 23 pathways significantly associated with survival in both CNV and protein data. Here are the top-10 most significant pathways (sorted by protein data significance):

```
SigBoth_df
#> # A tibble: 23 x 4
#>    terms  description                                      rawp_PROT rawp_CN
V
#>    <chr>  <chr>                                                <dbl>    <dbl
>
#>  1 WP195  IL-1 signaling pathway                             0.00247 0.0222
#>  2 WP363  Wnt Signaling Pathway                              0.00485 0.0486
#>  3 WP2036 TNF related weak inducer of apoptosis (TWEAK)~   0.00641 0.0226
#>  4 WP3850 Factors and pathways affecting insulin-like g~   0.00700 0.0366
#>  5 WP2203 Thymic Stromal LymphoPoietin (TSLP) Signaling~   0.0156  0.0181
#>  6 WP4136 Fibrin Complement Receptor 3 Signaling Pathway   0.0160  0.00035
8
#>  7 WP384  Apoptosis Modulation by HSP70                      0.0165  0.0311
#>  8 WP2018 RANKL/RANK (Receptor activator of NFKB (ligan~   0.0170  0.00041
8
#>  9 WP560  TGF-beta Receptor Signaling                        0.0205  0.0355
#> 10 WP2380 Brain-Derived Neurotrophic Factor (BDNF) sign~   0.0206  0.0113
#> # ... with 13 more rows
```

Similar to the protein pathway analysis shown in Section 2.3.4, we can also identify relevant genes with nonzero loadings that drives pathway significance in CNV. The “IL-1 signaling pathway” (WP195) is significant in both CNV and protein data.

```
# Copy Number Loadings
CNVwp195_ls <- getPathPCLs(ovCNV_aespcOut, "WP195")
CNV195load_df <-
  CNVwp195_ls$Loadings %>%
```

```
  filter(abs(PC1) > 0) %>%
  rename(PC1_CNV = PC1)

# Protein Loadings
PROTwp195_ls <- getPathPCLs(ovarian_aespcOut, "WP195")
PROT195load_df <-
  PROTwp195_ls$Loadings %>%
  filter(abs(PC1) > 0) %>%
  rename(PC1_PROT = PC1)

# Intersection
inner_join(CNV195load_df, PROT195load_df)
#> Joining, by = "featureID"
#> # A tibble: 1 x 3
#>   featureID PC1_CNV PC1_PROT
#>   <chr>       <dbl>    <dbl>
#> 1 NFKB1       0.113    0.483
```

The result showed the NFKB1 gene was selected by AESPCA when testing IL-1 signaling pathway (WP195) with survival outcome in both CNV and protein pathway analysis.

Because the proteomics data were recorded on a subset of the subjects shown in the copy number data, we can further examine the relationship between CNV and protein expressions for this gene.

```
NFKB1data_df <- inner_join(
  copyNumberClean_df %>%
    select(Sample, NFKB1) %>%
    rename(CNV_NFKB1 = NFKB1),
  ovSurv_df %>%
    select(Sample, NFKB1) %>%
    rename(PROT_NFKB1 = NFKB1)
)
#> Joining, by = "Sample"
```

This Pearson test shows that the correlation between CNV and protein expression is highly significant for this gene.

```
NFKB1_cor <- cor.test(NFKB1data_df$CNV_NFKB1, NFKB1data_df$PROT_NFKB1)
NFKB1_cor

    Pearson's product-moment correlation

data:  NFKB1data_df$CNV_NFKB1 and NFKB1data_df$PROT_NFKB1
t = 4.8873, df = 81, p-value = 5.081e-06
alternative hypothesis: true correlation is not equal to 0
95 percent confidence interval:
 0.2915318 0.6282385
sample estimates:
```

```
      cor
0.4772138
```

We can then visualize the multi-omics relationship for the NFKB1 gene using a scatterplot.

```
ggplot(data = NFKB1data_df) +
  # Set overall appearance
  theme_bw() +
  # Define the dependent and independent variables
  aes(x = CNV_NFKB1, y = PROT_NFKB1) +
  # Add the scatterplot
  geom_point(size = 2) +
  # Add a trendline
  geom_smooth(method = lm, se = FALSE, size = 1) +
  # Set main and axis titles
  labs(
    title = "NFKB1 Expressions in Multi-Omics Data",
    x = "Copy Number Variation",
    y = "Protein Expression"
  ) +
  # Include the correlation on the graph
  annotate(
    geom = "text", x = 0.5, y = -0.6,
    label = paste("rho = ", round(NFKB1_cor$estimate, 3))
  ) +
  annotate(
    geom = "text", x = 0.5, y = -0.7,
    label = "p-val < 10^-5"
  )
```

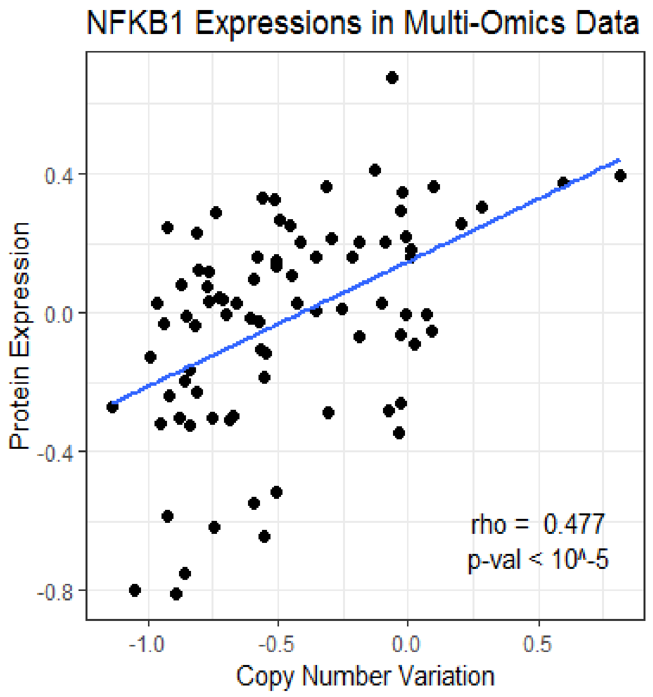

#### 3.3 An integrative view on patient-specific pathway activities

In Section 2.3.5, we have seen that there can be considerable heterogeneity in pathway activities between the patients. One possible reason could be that copy number changes might not result in changes in protein expression for some of the patients. `pathwayPCA` can be used to estimate pathway activities for each patient, both for protein expressions and copy number expressions separately. These estimates can then be viewed jointly using a Circos plot.

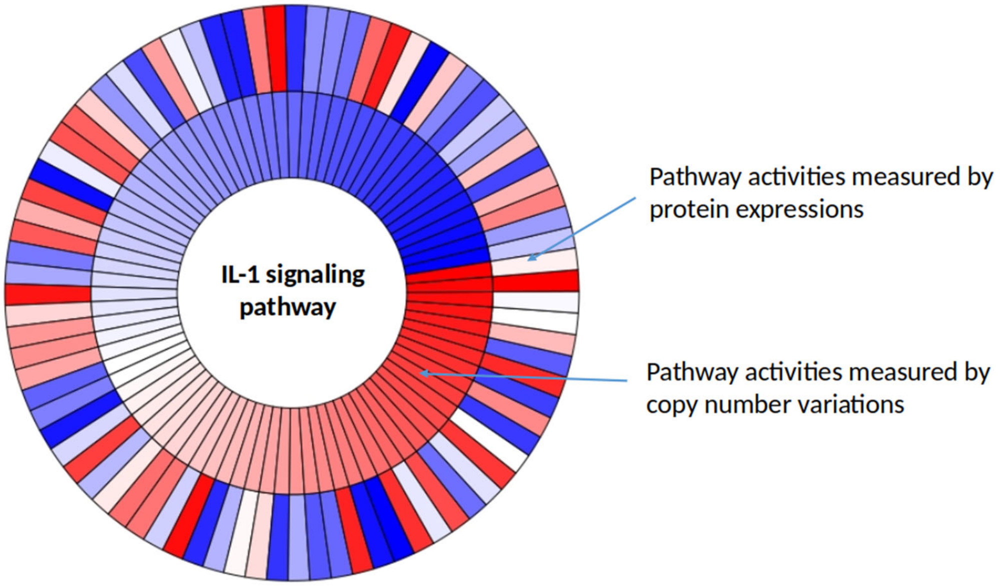

The accompanying Circos plot shown normalized copy number (outer circle) and protein expression (inner circle) pathway activities for the IL-1 signaling pathway in the ovarian cancer dataset samples. Each bar corresponds to a patient sample. Red bars indicate higher expression values and more pathway activity for the sample, while blue color bars indicate lower expression values and lower pathway activity for the sample. Note that only some patients have concordant changes in copy number and protein expression.

### 4. Case study: analysis of studies with complex designs

`pathwayPCA` is capable of analyzing studies with multiple experimental factors. In this section, we illustrate using `pathwayPCA` to test differential association of pathway expression with survival outcome in male and female subjects.

#### 4.1 Data setup and AESPCA analysis

For this example, we used TCGA KIRP RNAseq dataset downloaded from Xena datahub. First, we load the KIRP data, create an `Omics` data container, and extract first AESPC from each pathway.

```
# Load Data
kidney_df <- readRDS(paste0(dataDir_path, "/KIRP_Surv_RNAseq_inner.RDS"))

# Create Omics Container
kidney_Omics <- CreateOmics(
  assayData_df = kidney_df[, -(2:4)],
  pathwayCollection_ls = wikipathways_PC,
  response = kidney_df[, 1:3],
  respType = "surv",
  minPathSize = 5
)

  ======  Creating object of class OmicsSurv  =======
The input pathway database included 5831 unique features.
The input assay dataset included 20211 features.
Only pathways with at least 5 or more features included in the assay dataset are
  tested (specified by minPathSize parameter). There are 423 pathways which meet
  this criterion.
Because pathwayPCA is a self-contained test (PMID: 17303618), only features in
  both assay data and pathway database are considered for analysis. There are 5566
  such features shared by the input assay and pathway database.

# AESPCA
kidney_aespcOut <- AESPCA_pVals(
  object = kidney_Omics,
  numPCs = 1,
  parallel = TRUE,
  numCores = 2,
  numReps = 0,
  adjustment = "BH"
)

Part 1: Calculate Pathway AES-PCs
Initializing Computing Cluster: DONE
Extracting Pathway PCs in Parallel: DONE

Part 2: Calculate Pathway p-Values
Initializing Computing Cluster: DONE
Extracting Pathway p-Values in Parallel: DONE

Part 3: Adjusting p-Values and Sorting Pathway p-Value Data Frame
DONE
```

#### 4.2 Test for sex interaction with first PC

Next we can test whether there is differential pathway association with survival outcome for males and females by the following model,

$$h(t) = h_0(t)\exp[\beta_1 \mathrm{PC}_1 + \beta_2 \mathrm{male} + \beta_3(\mathrm{PC}_1 \times \mathrm{male})].$$

In this model, $h(t)$ is expected hazard at time $t$, $h_0(t)$ is baseline hazard when all predictors are zero, variable $male$ is an indicator variable for male samples, and $PC_1$ is a pathway's estimated first principal component based on AESPCA.

In order to test the sex interaction effect for all pathways, we will write a function which tests the interaction effect for one pathway.

```
TestIntxn <- function(pathway, pcaOut, resp_df){

  # For the given pathway, extract the PCs and loadings from the pcaOut list
  PCL_ls <- getPathPCLs(pcaOut, pathway)
  # Select and rename the PC
  PC_df <- PCL_ls$PCs %>% select(PC1 = V1)
  # Bind this PC to the phenotype data
  data_df <- bind_cols(resp_df, PC_df)

  # Construct a survival model with sex interaction
  sex_mod <- coxph(
    Surv(time, status) ~ PC1 + male + PC1 * male, data = data_df
  )
  # Extract the model fit statistics for the interaction
  modStats_mat <- t(
    coef(summary(sex_mod))["PC1:maleTRUE", ]
  )

  # Return a data frame of the pathway and model statistics
  list(
    statistics = data.frame(
      terms = pathway,
      description = PCL_ls$description,
      modStats_mat,
      stringsAsFactors = FALSE
    ),
    model = sex_mod
  )

}
```

As an example, we can test it on patwhay WP195,

```
TestIntxn("WP195", kidney_aespcOut, kidney_df[, 2:4])$model
#> Call:
#> coxph(formula = Surv(time, status) ~ PC1 + male + PC1 * male,
#>     data = data_df)
#>
#>                  coef exp(coef) se(coef)      z     p
#> PC1           0.03278   1.03332  0.08206  0.399 0.690
#> maleTRUE     -0.38822   0.67826  0.32827 -1.183 0.237
#> PC1:maleTRUE  0.05649   1.05812  0.10438  0.541 0.588
#>
```

```
#> Likelihood ratio test=3.97  on 3 df, p=0.2652
#> n= 320, number of events= 51
```

We can also apply this function to a list of pathways and select the significant pathways:

```
paths_char <- kidney_aespcOut$pVals_df$terms
interactions_ls <- lapply(
    paths_char,
    FUN = TestIntxn,
    pcaOut = kidney_aespcOut,
    resp_df = kidney_df[, 2:4]
)
names(interactions_ls) <- paths_char

interactions_df <-
  # Take list of interactions
  interactions_ls %>%
  # select the first element (the data frame of model stats)
  lapply(`[[`, 1) %>%
  # stack these data frames
  bind_rows() %>%
  as.tibble() %>%
  # sort the rows by significance
  arrange(`Pr...z..`)
#> Warning: `as.tibble()` is deprecated, use `as_tibble()` (but mind the new
semantics).
#> This warning is displayed once per session.

interactions_df %>%
  filter(`Pr...z..` < 0.05)
#> # A tibble: 14 x 7
#>    terms  description                  coef exp.coef. se.coef.     z Pr...z.
.
#>    <chr>  <chr>                       <dbl>     <dbl>    <dbl> <dbl>    <dbl
>
#>  1 WP1559 TFs Regulate miRNAs rel~ -0.597     0.550    0.216 -2.76  0.0057
3
#>  2 WP453  Inflammatory Response P~ -0.272     0.762    0.109 -2.50  0.0126
#>  3 WP3893 Development and heterog~ -0.329     0.719    0.137 -2.40  0.0166
#>  4 WP3929 Chemokine signaling pat~ -0.252     0.777    0.109 -2.32  0.0206
#>  5 WP2849 Hematopoietic Stem Cell~ -0.278     0.757    0.129 -2.16  0.0305
#>  6 WP1423 Ganglio Sphingolipid Me~ -0.487     0.615    0.230 -2.11  0.0346
#>  7 WP3892 Development of pulmonar~ -0.316     0.729    0.151 -2.10  0.0360
#>  8 WP3678 Amplification and Expan~ -0.357     0.700    0.171 -2.09  0.0363
#>  9 WP4141 PI3K/AKT/mTOR - VitD3 S~  0.314     1.37     0.153  2.05  0.0407
#> 10 WP3941 Oxidative Damage         -0.354     0.702    0.176 -2.01  0.0442
#> 11 WP3863 T-Cell antigen Receptor~ -0.233     0.792    0.117 -1.99  0.0462
#> 12 WP3872 Regulation of Apoptosis~  0.267     1.31     0.135  1.98  0.0472
#> 13 WP3672 LncRNA-mediated mechani~ -0.392     0.675    0.198 -1.98  0.0475
#> 14 WP3967 miR-509-3p alteration o~ -0.299     0.741    0.151 -1.98  0.0481
```

The results showed the most significant pathway is WP1559 (“TFs Regulate miRNAs related to cardiac hypertrophy”). We can inspect the model results for this pathway directly.

```
summary(interactions_ls[["WP1559"]]$model)
#> Call:
#> coxph(formula = Surv(time, status) ~ PC1 + male + PC1 * male,
#>     data = data_df)
#>
#>   n= 320, number of events= 51
#>
#>                  coef exp(coef) se(coef)      z Pr(>|z|)
#> PC1            0.7534    2.1242   0.1782  4.228 2.36e-05 ***
#> maleTRUE       0.1825    1.2002   0.4268  0.427  0.66903
#> PC1:maleTRUE -0.5974    0.5502   0.2162 -2.763  0.00573 **
#> ---
#> Signif. codes:  0 '***' 0.001 '**' 0.01 '*' 0.05 '.' 0.1 ' ' 1
#>
#>              exp(coef) exp(-coef) lower .95 upper .95
#> PC1             2.1242     0.4708    1.4980    3.0122
#> maleTRUE        1.2002     0.8332    0.5199    2.7705
#> PC1:maleTRUE    0.5502     1.8174    0.3602    0.8406
#>
#> Concordance= 0.654  (se = 0.047 )
#> Rsquare= 0.068   (max possible= 0.792 )
#> Likelihood ratio test= 22.51  on 3 df,   p=5e-05
#> Wald test            = 30.03  on 3 df,   p=1e-06
#> Score (logrank) test = 33.09  on 3 df,   p=3e-07
```

The results showed that although sex is not significantly associated with survival outcome, the association of pathway gene expression (PC1) with survival is highly dependent on sex of the samples.

#### 4.3 Survival curves by sex interaction

For visualization, we can divide subjects according to sex and high or low PC-expression, and add a group indicator for the four groups.

```
# Bind the Pheno Data to WP1559's First PC
kidneyWP1559_df <- bind_cols(
  kidney_df[, 2:4],
  getPathPCLs(kidney_aespcOut, "WP1559")$PCs %>% select(PC1 = V1)
)

# Add Grouping Feature
kidneySurvWP1559grouped_df <-
  kidneyWP1559_df %>%
  # add strength indicator
  mutate(direction = ifelse(PC1 > median(PC1), "High", "Low")) %>%
  # group by interaction of sex and strength on PC
```

```
  mutate(group = paste0(direction, ifelse(male, "_Male", "_Female"))) %>%
  # remove summarized columns
  select(-male, -PC1, -direction)
```

Now we can plot survival curves for the four groups.

```
# Fit the survival model
fit <- survfit(
  Surv(time, status) ~ group,
  data = kidneySurvWP1559grouped_df
)

ggsurvplot(
  fit, conf.int = FALSE,
  xlab = "Time in Days",
  ylim = c(0.4, 1)
)
```

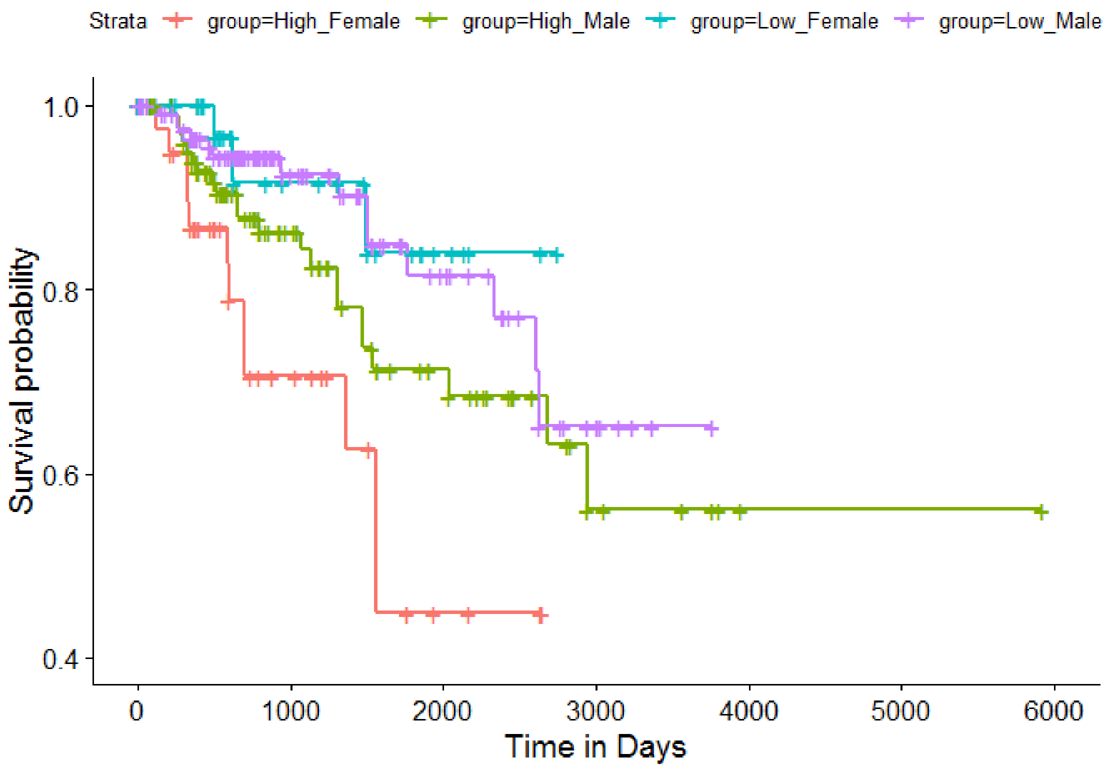

These Kaplan-Meier curves showed that while high or low pathway activities were not associated with survival in male subjects (green and purple curves, respectively), female subjects with high pathway activities (red) had significantly worse survival outcomes than those with low pathway activities (blue).

#### 5. Further reading

For addtional information on `pathwayPCA`, please see each of our supplementary vignette chapters for detailed tutorials on each of the three topics discussed above. These vignettes are:

- Chapter 2: Import Data
- Chapter 3: Create Omics Data Objects
- Chapter 4: Test Pathway Significance
- Chapter 5: Visualizing the Results

#### 6. References

Chen, X., Wang, L., Smith, J.D. and Zhang, B. (2008) Supervised principal component analysis for gene set enrichment of microarray data with continuous or survival outcomes. Bioinformatics, 24, 2474-2481.

Chen, X., Wang, L., Hu, B., Guo, M., Barnard, J. and Zhu, X. (2010) Pathway-based analysis for genome-wide association studies using supervised principal components. Genetic epidemiology, 34, 716-724.

Chen, X. (2011) Adaptive elastic-net sparse principal component analysis for pathway association testing. Statistical applications in genetics and molecular biology, 10.

The R session information for this vignette is:

```
sessionInfo()
#> R version 3.5.2 (2018-12-20)
#> Platform: x86_64-w64-mingw32/x64 (64-bit)
#> Running under: Windows 7 x64 (build 7601) Service Pack 1
#>
#> Matrix products: default
#>
#> locale:
#> [1] LC_COLLATE=English_United States.1252
#> [2] LC_CTYPE=English_United States.1252
#> [3] LC_MONETARY=English_United States.1252
#> [4] LC_NUMERIC=C
#> [5] LC_TIME=English_United States.1252
#>
#> attached base packages:
#> [1] stats     graphics  grDevices utils     datasets  methods   base
#>
#> other attached packages:
#>  [1] survminer_0.4.3   ggpubr_0.2        magrittr_1.5
#>  [4] survival_2.43-3   bindrcpp_0.2.2    pathwayPCA_0.99.1
#>  [7] forcats_0.3.0     stringr_1.3.1     dplyr_0.7.8
#> [10] purrr_0.3.0       readr_1.3.1       tidyr_0.8.2
```

```
#> [13] tibble_2.0.1      ggplot2_3.1.0     tidyverse_1.2.1
#> 
#> loaded via a namespace (and not attached):
#>  [1] Rcpp_1.0.0        lubridate_1.7.4   lattice_0.20-38  
#>  [4] zoo_1.8-4         assertthat_0.2.0  digest_0.6.18    
#>  [7] utf8_1.1.4        R6_2.3.0          cellranger_1.1.0 
#> [10] plyr_1.8.4        backports_1.1.3   lars_1.2         
#> [13] evaluate_0.12     httr_1.4.0        pillar_1.3.1     
#> [16] rlang_0.3.1       lazyeval_0.2.1    readxl_1.2.0     
#> [19] data.table_1.12.0 rstudioapi_0.9.0  Matrix_1.2-15    
#> [22] rmarkdown_1.11    labeling_0.3      splines_3.5.2    
#> [25] munsell_0.5.0     broom_0.5.1       compiler_3.5.2   
#> [28] modelr_0.1.2      xfun_0.4          pkgconfig_2.0.2  
#> [31] htmltools_0.3.6   tidyselect_0.2.5  gridExtra_2.3    
#> [34] km.ci_0.5-2       fansi_0.4.0       crayon_1.3.4     
#> [37] withr_2.1.2       grid_3.5.2        xtable_1.8-3     
#> [40] nlme_3.1-137      jsonlite_1.6      gtable_0.2.0     
#> [43] KMsurv_0.1-5      scales_1.0.0      cli_1.0.1        
#> [46] stringi_1.2.4     xml2_1.2.0        survMisc_0.5.5   
#> [49] generics_0.0.2    tools_3.5.2       cmprsk_2.2-7     
#> [52] glue_1.3.0        hms_0.4.2         parallel_3.5.2   
#> [55] yaml_2.2.0        colorspace_1.4-0  rvest_0.3.2      
#> [58] knitr_1.21        bindr_0.1.1       haven_2.0.0
```
